# Contrasting drugs from decoys

**DOI:** 10.1101/2022.11.03.515086

**Authors:** Samuel Sledzieski, Rohit Singh, Lenore Cowen, Bonnie Berger

## Abstract

Protein language models (PLMs) have recently been proposed to advance drugtarget interaction (DTI) prediction, and have shown state-of-the-art performance on several standard benchmarks. However, a remaining challenge for all DTI prediction models (including PLM-based ones) is distinguishing true drugs from highly-similar decoys. Leveraging techniques from self-supervised contrastive learning, we introduce a second-generation PLM-based DTI model trained on triplets of proteins, drugs, and decoys (small drug-like molecules that do not bind to the protein). We show that our approach, **CON-Plex**, improves specificity while maintaining high prediction accuracy and generalizability to new drug classes. CON-Plex maps proteins and drugs to a shared latent space which can be interpreted to identify mutually-compatible classes of proteins and drugs. Data and code are available at https://zenodo.org/record/7127229.

## 1 Introduction

Accurate and rapid prediction of drug-target interactions (DTIs) remains a major open problem in drug discovery. A DTI prediction method needs to address the twin goals of generalizability and specificity. The ideal predictive approach would generalize broadly, i.e., make accurate predictions on hitherto unseen classes of drugs. It would also have fine-grained specificity, i.e., be able to discern the binding impact of minor changes to a protein or drug molecule. Even in early DTI prediction methods, it was observed that these goals are difficult to achieve simultaneously. In 2006, Huang et al. introduced the Directory of Useful Decoys (DUD) [9] (later updated to DUD-E, “DUD Enhanced” [16]) to enable quantification of prediction specificity. For a set of DTIs, DUD-E offers *decoys:* small molecules that share physical and chemical similarities with the true-positive drug but do not actually bind with the target. Huang et al.’s work presaged the development of similar ideas in adversarial machine learning. In particular, their formulation of the DTI prediction task is similar to generative adversarial networks (GANs) [7]: the role of a generator is played by the DUD-E database, while the DTI predictor acts as a discriminator.

In this paper, we seek to substantially improve the specificity of DTI prediction while retaining high accuracy and generalizability. We approach the DTI prediction task from the perspective of protein language models (PLMs). Trained over hundreds of millions of protein sequences, PLMs apply the distributional hypothesis [2, 3, 18, 4] and learn a rich implicit featurization of proteins that has proven useful in a variety of tasks [12, 19, 21]. Sledzieski et al. [20], and later Goldman et al. [6] independently, have demonstrated the power of PLMs for DTI prediction. The use of pre-trained PLMs unlocks the full richness and diversity of data across the protein universe, whereas models trained solely on DTI data can leverage only the very limited percentage of protein space that has been tested experimentally for interactions.

Here, we present **CON-Plex**, a sequence-based DTI prediction approach that builds on previous PLM-based methods. The key conceptual advance of this paper is the adaptation of mechanisms for self-supervised contrastive learning to enable discrimination of drugs from decoys. Anchored by the lexicographic protein representation from a pre-trained PLM, we show that this contrastive approach enables CON-Plex to retain high predictive accuracy.

The triplet loss function, commonly used in contrastive self-supervised learning [1], compares three entities of the same type: an anchor, a positive example (sometimes generated by modifying the anchor itself), and a negative example. However, it is not immediately clear how this loss function should be employed in the DTI setting, as the latter corresponds to a supervised learning task and involves entities of two different types (i.e., protein and small molecule). Our key insight is that using a formulation where these entities are mapped to a shared latent space allows the triplet loss to be applied (with the protein, drug and a decoy as the triplet). CON-Plex introduces a co-training framework that utilizes a ground-truth DTI dataset alongside the DUD-E database and modifies the standard triplet loss to incorporate a margin annealing scheme.

We show that the CON-Plex approach substantially improves the ability to distinguish test-set drugs from decoys while retaining high DTI prediction accuracy. Furthermore, it also makes the shared latent space of drugs and proteins more robust and interpretable, and we show that the learned representations of drugs cluster better and are closer to the target after contrastive training.

## 2 Methods

### Data

We trained and evaluated CON-Plex on examples from the Directory of Useful Decoys: Enhanced (**DUD-E**) [16]. We simultaneously co-trained the model also on the DTIs from the **BIOSNAP** [22] benchmark. Statistics for each data set, including number of unique proteins and drugs, and number of training/validation/test edges are provided in Table A.1.

DUD-E consists of 102 proteins and 22,886 known drugs (average 224 molecules per target). For each drug, 50 decoys are available. We focused on the 57 targets (and their associated drugs/decoys) where the enzyme category was specified; these spanned GPCRs, kinases, nuclear proteins, and proteases. We stratified train-test splits by categories, so that there are targets of each category in both the training and test sets. We provide the full list of targets in Table A.2.

Our inclusion of the BIOSNAP dataset was motivated by the relatively few ground-truth DTIs contained in DUD-E, which are insufficient to learn drug-target binding. However, the BIOSNAP benchmark consists of only positive DTIs. Following Huang et al. [8], we made the assumption that a random pair is unlikely to be positively interacting and created negative DTIs by randomly sampling an equal number of protein-drug pairs. We train on a random 70% split, leaving 10% for validation, and the remaining 20% for testing.

### Sledzieski et al.’s PLM-Based DTI Model

CON-Plex is based on the work of Sledzieski et al. [20], which exploits a highly-informative PLM representation of proteins to predict DTIs. We briefly describe the base model (**PLM-DTI**) here, which we then augment with the contrastive training. Sledzieski et al. featurize the protein target using a pre-trained ProtBert model [4]. Given a protein sequence, ProtBert generates an embedding *T_full_* ∈ ℝ^*n×d_t_*^ (*d* = 1024) for a protein of length *n*, which is then mean-pooled along the length of the protein, resulting in a vector *T* ∈ ℝ^*d_t_*^. Simultaneously, they generate a feature vector for the drug using its Morgan fingerprint [15], a fixed-length embedding *M* ∈ ℝ^*d_m_*^ (*d_m_* = 2048) of the drug’s SMILES string produced by considering the local neighborhood around atoms in the molecular graph.

Given the embeddings of target *T* ∈ ℝ^*dt*^ and small molecule *M* ∈ ℝ^*dm*^, Sledzieski et al. transform them into *T*\* = *FC*(*T*), *M*\* = *FC*(*T*) ∈ ℝ^*d*^, where *FC*(·) is a single fully-connected layer with a ReLU activation. The latent space embeddings *T*\*, *M*\* are then used to compute the probability of a drug-target interaction 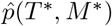 as the cosine similarity between the embedding vectors, followed by a sigmoid activation.

### Contrastive Loss

We build upon Sledzieski et al.’s model architecture, training CON-Plex in alternating epochs, one focused on the BIOSNAP dataset and the other on the DUD-E decoy dataset. On the BIOSNAP data, model weights are updated using the binary cross-entropy loss (*L_BCE_*). In alternate epochs, the model is supervised on decoys, with model weights updated using an adaptation of the triplet distance loss. The conventional use of triplet loss in contrastive loss considers three training points of the same type, the **anchor**, **positive**, and **negative**, and aims to minimize the distance between the anchor and positive examples while maximizing the distance between the anchor and the negative examples.

In the DTI setting, the natural choice for a triplet is the protein target as the anchor, the true drug as the positive and decoy as the negative example, respectively. We derive a training set of triplets in the following manner: for each known interacting drug-target pair (*T, M*^+^), we randomly sample *k* = 50 non-interacting pairs (*T, M*^-^) and generate the triplets (*T, M*^+^, *M*^-^), where *M*^-^ is drawn from the set of all decoys against T. We map these to latent space embeddings as described above. Since all the entities are now comparable to each other, we can compute the triplet margin-distance loss (*L_TRM_*).

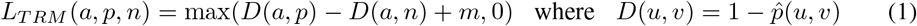

### Margin Annealing

The margin *m* sets the maximum required delta between distances, above which the loss is zero. Initially, a large margin requires the decoy to be much further from the target than the drug to avoid a penalty, resulting in larger weight updates. As training progresses, lower margins relax this constraint, requiring only that the drug be closer than the decoy as *m* → 0. Here, the margin is initialized at *M_max_* = 0.25 and decreased over *E_max_* = 50 contrastive epochs according to a tanh decay schedule. At epoch *i*, the margin is set to

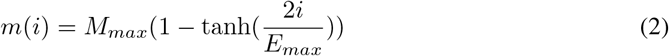

### Implementation

Model weights were initialized using the Xavier method from a normal distribution [5] and optimized with the AdamW [14] optimizer for 50 BCE epochs and 50 contrastive epochs, interleaved. For BIOSNAP (DUD-E) training, the learning rate was initially set to 10^-4^ (10^-5^) and adjusted according to a cosine annealing schedule with warm restarts [13] every 10 epochs. We used a latent dimension *d* = 1024 (results were robust to variations in latent dimension size) and a batch size of 32. The model was implemented in PyTorch version 1.11 on a single NVIDIA A100 GPU; after data loading, the full training run completed in ~20-25 mins.

## 3 Results

### Decoy Discrimination

We demonstrate that CON-Plex achieves extremely high performance on the decoy discrimination task, out-performing two baseline models trained solely on binary drug-target or decoy-target pairs, a logistic regression (**LR**) and multi-layer perceptron (**MLP**) [17]. The MLP was initialized with a single hidden layer with 1024 nodes, and thus has the same number of parameters as CON-Plex (3,147,776). We report the area under precision-recall curve (AUPR) and receiver operating characterstic curve (AUROC) in Table 1. The baseline models have AUPRs of 0.257 (LR) and 0.306 (MLP), while CON-Plex has an AUPR of 0.453. We also report the performance of Sledzieski et al.’s PLM-based DTI model trained only on BIOSNAP (**PLM-DTI**). While this model performs well on general DTI prediction, it is unable to discriminate between the highly-similar decoy drugs and only reaches an AUPR of 0.076—demonstrating the need for contrastive co-training.

**Table 1:**
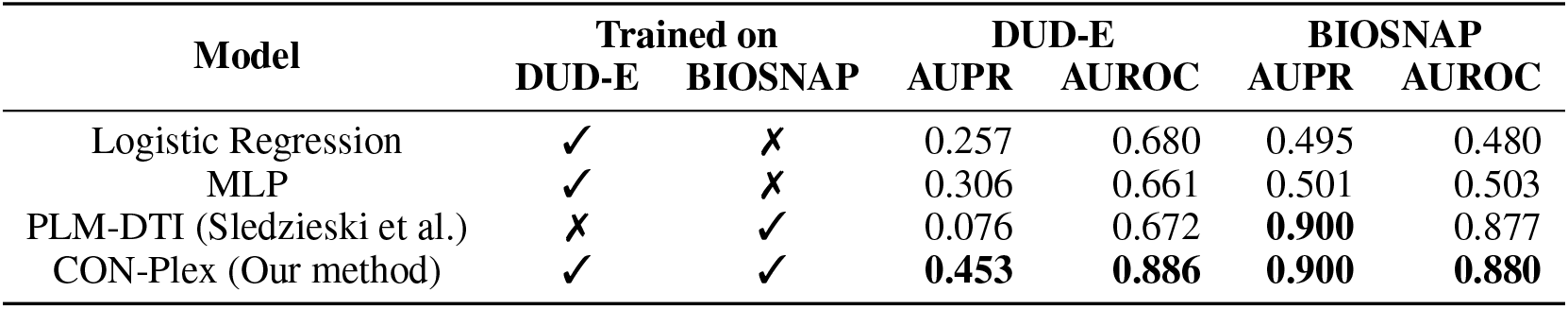
CON-Plex achieves state-of-the-art performance on decoy detection and DTI prediction. We report the AUPR and AUROC for each method on DUD-E, a decoy determination benchmark, and on BIOSNAP, a general DTI benchmark. CON-Plex, which was trained on both BIOSNAP and DUDE, outperforms two baseline models (Logistic Regression, MLP) that were trained specifically to perform decoy discrimination. We show that co-training on decoys does not hurt general performance, as CON-Plex matches PLM-DTI (Sledzieski et al. [20]) trained only on BIOSNAP.

| Model | Trained on |  | DUD-E |  | BIOSNAP |  |
| --- | --- | --- | --- | --- | --- | --- |
|  | DUD-E | BIOSNAP | AUPR | AUROC | AUPR | AUROC |
| Logistic Regression | ✓ | ✗ | 0.257 | 0.680 | 0.495 | 0.480 |
| MLP | ✓ | ✗ | 0.306 | 0.661 | 0.501 | 0.503 |
| PLM-DTI (Sledzieski et al.) | ✗ | ✓ | 0.076 | 0.672 | **0.900** | 0.877 |
| CON-Plex (Our method) | ✓ | ✓ | **0.453** | **0.886** | **0.900** | **0.880** |

### General DTI Performance

While contrastive training has an extremely beneficial impact on decoy discrimination, it is desirable that this additional training does not decrease overall DTI performance. On the BIOSNAP test set, CON-Plex has an equivalent AUPR to the non-contrastive PLM-DTI (Table 1). Unsurprisingly, the baseline models trained only on decoy discrimination are unable to predict DTIs and show essentially random performance on the BIOSNAP data.

### Learning meaningful representations of ligand binding

The co-embedding approach taken by PLM-DTI and CON-Plex also enables interpretability— we can visualize the shared latent space, and measure how the representations of proteins and drugs change as a result of the contrastive training. For each model and each target in the DUD-E test set, we plotted the target alongside all drugs and decoys using the embeddings from the base and contrastive-tuned model. Figure 2a,b shows one such example, the tyrosine kinase *VGFR2*. We also show the distribution of distances in the latent space between the target embedding and the embeddings of the drugs and decoys for each model (Figure 2c, d) (p-values from one-sided t-test).

**Figure 1:**
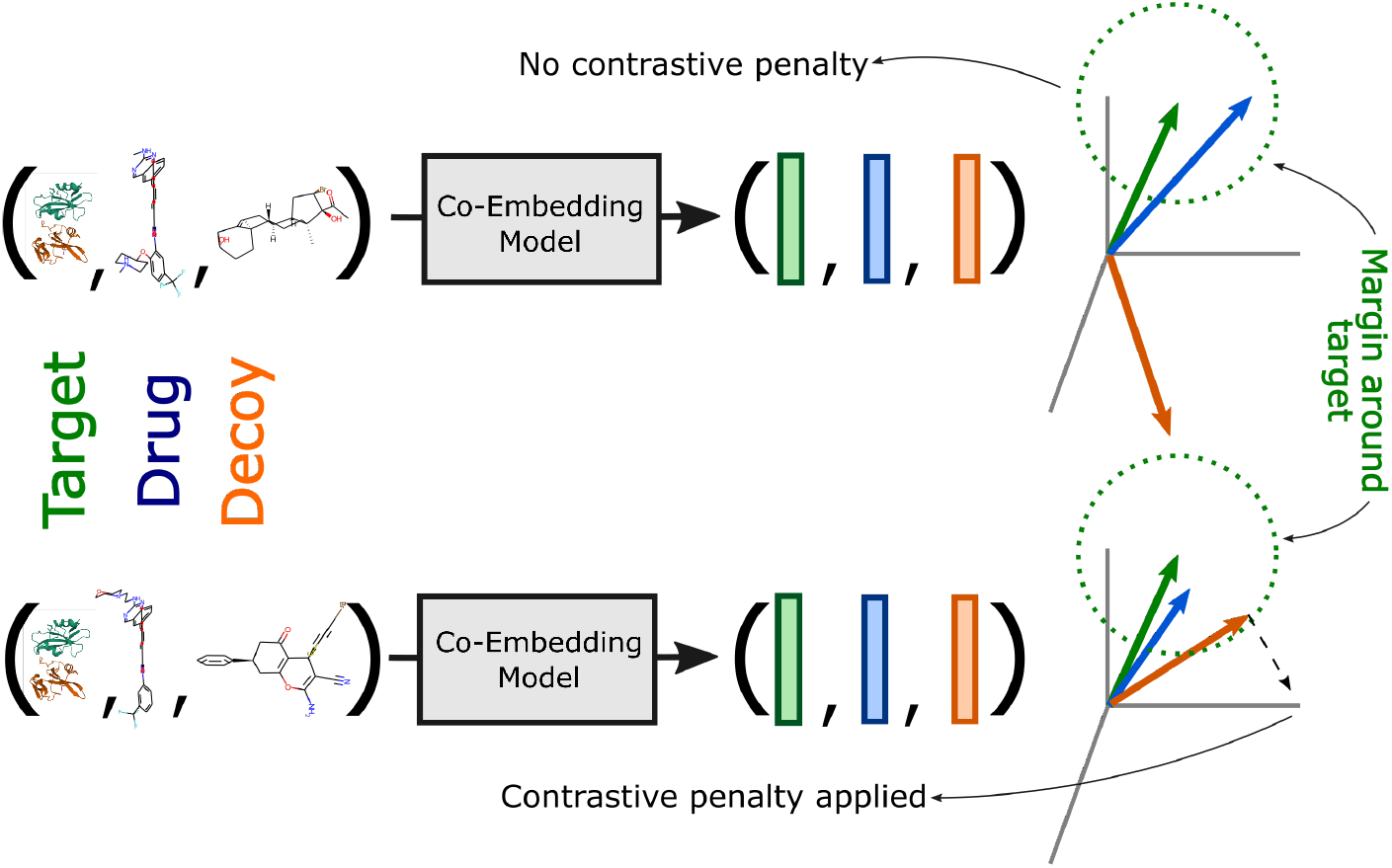
Overview of contrastive learning in CON-Plex. We augment the PLM-based DTI model introduced by Sledzieski et al. [20] with contrastive learning to achieve highly specific decoy detection. Given a triplet of a protein target, a small molecule that binds to it (“drug”), and a small molecule that doesn’t (“decoy”), we co-embed all of them into a shared latent space and apply a contrastive penalty if the decoy is closer (or further but within some margin) to the target than the drug (bottom). If the decoy is sufficiently far away from the target (top), no additional penalty is applied. Concurrently, the model is co-trained on a separate binary classification task, using a DTI data set that contains a greater diversity of unique proteins but lacks decoys.

**Figure 2:**
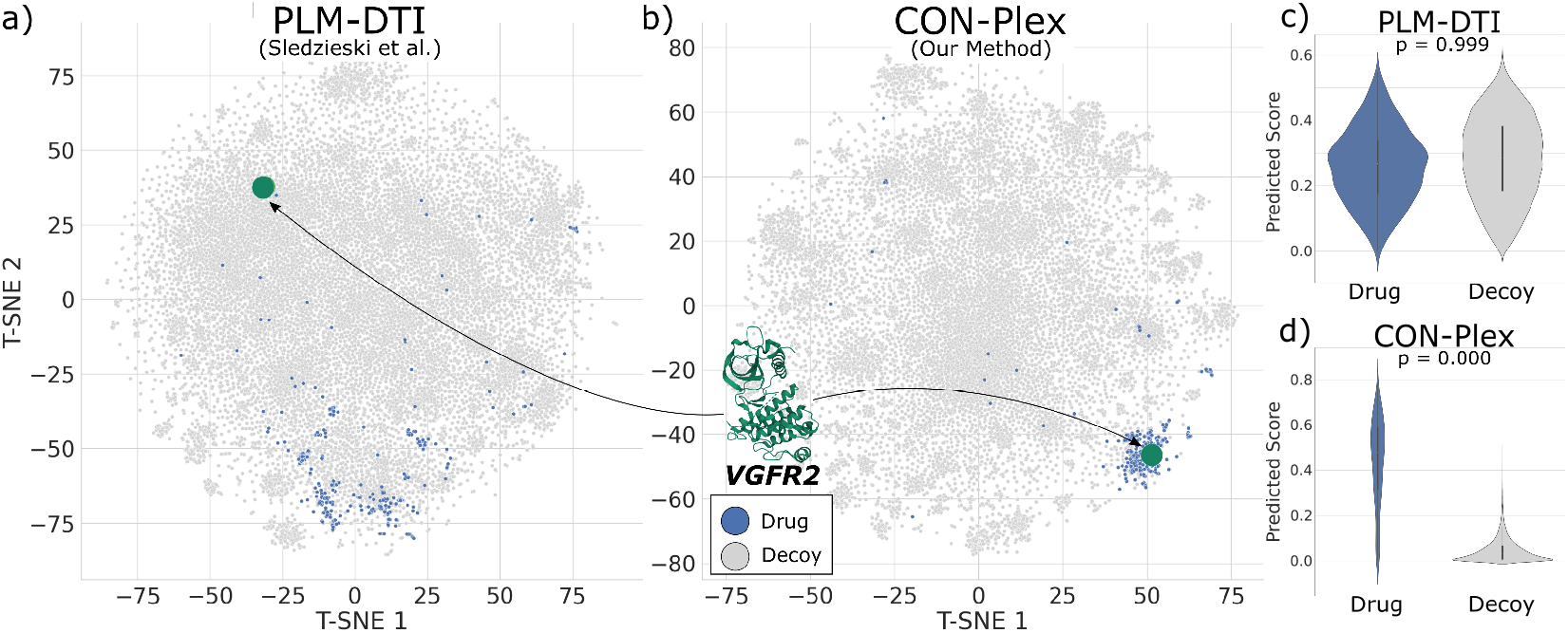
Improved latent space representation using contrastive learning. We compare the learned co-embedding of the baseline PLM-DTI (Sledzieski et al. [20]) with the shared latent space learned by CON-Plex. **(a)** In the original space, drugs (blue) of *VGFR2* are relatively scattered and are far from the target embedding (green). **(b)** In the CON-Plex space, drugs cluster around the target embedding. **(c)** Using PLM-DTI, it is difficult to distinguish drugs and decoys based on their distance to *VGFR2 (p* = 0.999, one-sided t-test). **(d)** CON-Plex clearly differentiates drugs and decoys (*p* = 0.000).

In Figure 3, we show a quantitative analysis of all test-set targets. We compute the effect size (Cohen’s *d*) of the difference between predicted drug and decoy scores. We plot these effect sizes for both PLM-DTI and CON-Plex. An increase in the effect size indicates that the co-embedding distances learned by the model better represent binding specificity. For each class of targets, we also report the median *p*-value (one-sided t-test) between drug and decoy scores predicted by CON-Plex. For all targets, contrastive training produced a stronger latent-space separation between drugs and decoys.

**Figure 3:**
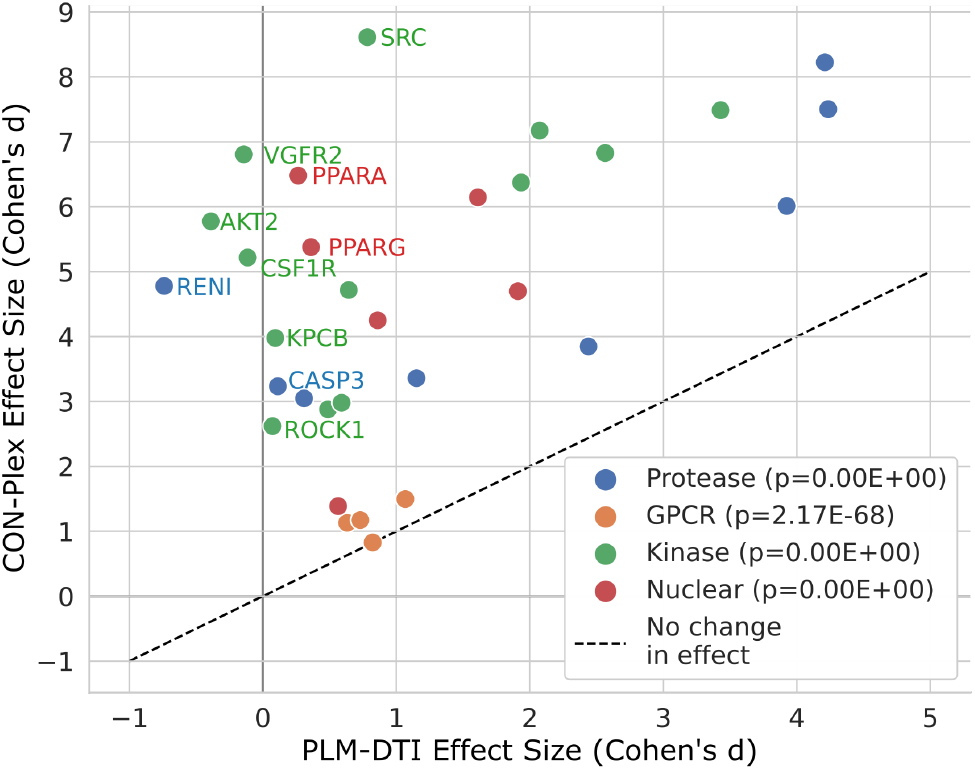
Contrastive training improves separation between drugs and decoys. Difference in predicted drug vs. decoy score is measured per target using effect size (Cohen’s d). For each class, we additionally report the median p-value (one-sided t-test) for CON-Plex drug vs. decoy scores. The *y* = *x* line corresponds to no change in effect as a result of contrastive training. The change in effect is strongest for kinases and nuclear proteins, while contrastive training has a weaker effect for GPCRs. Targets where effect size improves by at least 10x are labeled.

## 4 Discussion

High performing DTI prediction methods should be able to generalize broadly to unseen types of drugs and targets, while also discriminating between highly similar molecules with different binding properties. Previous work demonstrated the utility of PLMs to improve the generalizability of DTI prediction methods[6, 20]; we now add a contrastive learning approach which improves specificity. The contrastive approach taken by CON-Plex is directly enabled by the architecture of the base PLM-enabled lexicographic model— to compute the triplet distance loss, the protein and drugs must be co-embedded, and the distance between them must be meaningful and simply computed. Such an approach would not be feasible using a model which concatenates features up front, nor for a model which has significant computation (say, additional linear layers) defining the probability of interaction after the co-embedding. Thus the shared lexicographic space in which we embed the proteins, targets, and decoys is key. A limitation of CON-Plex, and of previous approaches, is that it only provides a binary output and not an affinity prediction or the structural mechanism of interaction. CON-Plex approaches the DTI decoy problem from the perspective of adversarial machine learning, where the model must act as a discriminator for adversarial examples from the decoy database. Future work could explore adapting molecular generation methods such as JT-VAE or HierG2G [10, 11] to directly act as a generator for decoys. Notably, high-specificity DTI prediction is also valuable beyond decoy detection— the greater specificity of inference can help improve personalized medicine or the modeling of drug effects against rare variants from under-represented populations.

## Acknowledgments and Disclosure of Funding

The authors thank Kapil Devkota, Tristan Bepler, and Tim Truong for helpful discussions. SS was supported by the National Science Foundation Graduate Research Fellowship under Grant No. 2141064. RS and BB were supported by the NIH grant R35GM141861. LC was supported by NSF CCF-1934553. The authors have no competing interests to disclose.

## A Appendix

## A.1 Data Details

**Table A.1:** Data set summary. CON-Plex is trained in alternating epochs on DTI data from BIOSNAP [22] and on decoy data from DUD-E [16]. Protein targets in DUD-E are separated by class into proteases, GPCRs, kinases, and nuclear proteins. We report here the number of unique drugs and proteins in each data set, as well as the number of training, validation, and testing pairs (positive/negative) available to the model.

| Data Set | Drugs | Proteins | # Training | # Validation | # Test |
| --- | --- | --- | --- | --- | --- |
| DUD-E |  |  | 8996 / 406208 | — | 11430 / 521132 |
| Protease | 286089 | 15 |  |  |  |
| GPCR | 99671 | 5 |  |  |  |
| Kinase | 315399 | 26 |  |  |  |
| Nuclear | 151133 | 11 |  |  |  |
| BIOSNAP | 4510 | 2181 | 9670 / 9568 | 1396 / 1352 | 2770 / 2727 |

**Table A.2:** DUD-E target classes and splits. Targets were randomly split within each class, so that there were examples of each class in the training and test set. There are a total of 57 targets, 26 in training and 31 in testing.

|  |  |  |
| --- | --- | --- |
| AA2AR | Gpcr | Train |
| ABL1 | Kinase | Train |
| AKT1 | Kinase | Train |
| CDK2 | Kinase | Train |
| EGFR | Kinase | Train |
| FAK1 | Kinase | Train |
| FGFR1 | Kinase | Train |
| IGF1R | Kinase | Train |
| JAK2 | Kinase | Train |
| KIT | Kinase | Train |
| MAPK2 | Kinase | Train |
| MK01 | Kinase | Train |
| PLK1 | Kinase | Train |
| TGFR1 | Kinase | Train |
| ANDR | Nuclear | Train |
| ESR2 | Nuclear | Train |
| MCR | Nuclear | Train |
| PPARD | Nuclear | Train |
| THB | Nuclear | Train |
| ACE | Protease | Train |
| BACE1 | Protease | Train |
| DPP4 | Protease | Train |
| FA7 | Protease | Train |
| HIVPR | Protease | Train |
| MMP13 | Protease | Train |
| TRYB1 | Protease | Train |
| ADRB1 | Gpcr | Test |
| ADRB2 | Gpcr | Test |
| CXCR4 | Gpcr | Test |
| DRD3 | Gpcr | Test |
| AKT2 | Kinase | Test |
| BRAF | Kinase | Test |
| CSF1R | Kinase | Test |
| KPCB | Kinase | Test |
| LCK | Kinase | Test |
| MET | Kinase | Test |
| MK10 | Kinase | Test |
| MK14 | Kinase | Test |
| MP2K1 | Kinase | Test |
| ROCK1 | Kinase | Test |
| SRC | Kinase | Test |
| VGFR2 | Kinase | Test |
| WEE1 | Kinase | Test |
| ESR1 | Nuclear | Test |
| GCR | Nuclear | Test |
| PPARA | Nuclear | Test |
| PPARG | Nuclear | Test |
| PRGR | Nuclear | Test |
| RXRA | Nuclear | Test |
| ADA17 | Protease | Test |
| CASP3 | Protease | Test |
| FA10 | Protease | Test |
| LKHA4 | Protease | Test |
| RENI | Protease | Test |
| THRB | Protease | Test |
| TRY1 | Protease | Test |
| UROK | Protease | Test |

